# Dynamics of a Lipid Vesicle across a Microfluidic Constriction: How does the Fluidity of the Encapsulation and its Micro-Environment Matter?

**DOI:** 10.1101/2023.05.03.539209

**Authors:** Tanoy Kahali, Devi Prasad Panigrahi, Suman Chakraborty

## Abstract

Constriction in the flow passage in the physiological circulatory system is central to the occurrence of several diseased conditions such as thrombosis and is also pivotal towards the understanding of several regulatory processes in the human microvasculature. It is, therefore, imperative to advance a mechanistic insight on the dynamics of the transiting cellular encapsulations in a physiologically-mimicking micro-confinement, with particular focus on deciphering the role of its mechano-physical properties. Here we bring out a quantitative depiction on the role of the membrane fluidity and the initial deflation (shape deviation from sphericity) of a lipid vesicle during its morphological transition from stretching to tumbling via rolling as it migrates across a microfluidic constriction. Based on our experimental observations as well as theoretical insights, we construct a regime map to elucidate the range of the key dimensionless parameters orchestrating the dynamic transition. Our results further bring out the role of the initial position of the lipid vesicle on its subsequent stretching dynamics, exhibiting characteristic nonlinearities and non-monotonic trends. In addition, our observations on the vesicle’s stretching dynamics emerge from mapping selectively with the viscosity contrast between the encapsulated and the suspending fluid medium, offering potential physiologically relevant cues on the impact of the aging of a cellular moiety on its deformability as it transits through a constricted path. Such mechanistic insights may potentially enable establishing quantitative correlations between the dynamical transition of a cellular encapsulation and its mechano-physical properties, which may in turn, have decisive implications in various states of health and disease while circulating across microvascular fluidic pathways.

**Impact Statement:** This study brings out a quantitative mechanistic insight into the dynamics of migrating lipid vesicles as they migrate through a constricted microfluidic passage. Having a direct similitude with the movement of red blood cells in human microvascular pathways, the resulting mapping between the initial shape and bending properties of the vesicle membrane with three distinct morphological transitions (stretching, rolling, and tumbling) provides cues for understanding the healthy and diseased states of the cells based on their morpho-dynamic features and establish exclusive connectivity of the same with the cell membrane as well as the cytoplasm properties, a paradigm that is currently non-existant. This, in turn, may lead to a novel mechanistic approach of label-free disease detection based on cellular imaging, for which the current understanding is mostly empirical rather than fundamental.

## 1. Introduction

Comprehending the fluid dynamics of human microcirculation and its implication on various aspects of health and diseases delves critically into understanding the morpho-dynamic evolution of different cellular encapsulations in physiologically relevant fluidic pathways(B.M. Koeppen and B. A. Stanton n.d.; C. openStax 2017). Whereas physiological matter such as red blood cells (RBCs) have their unique complexities that cannot be represented by mechanistic considerations alone, their model idealizations, such as lipid vesicles, have emerged to be enormously effective in offering valuable insights on their interactions with the surrounding fluidic media(Badr Kaoui 2009; Faivre 2006; Kantsler n.d.). These encapsulations, akin to cells, are liquid-filled globules made of phospholipids bilayer membrane, engulfing internal fluidic matter analogous to the cytoplasm (Kahali *et al*. 2022; Mader *et al*. 2006; Marella and Udaykumar 2004; Tran-Son-Tay *et al*. 1998). Despite a gross simplistic idealization, the dynamics of such model vesicles has been established to mimic quite closely the dynamics of RBCs and other similar active biological encapsulations, and thus has remained in the forefront of bio-fluid mechanics research over the last two decades (Abkarian *et al*. 2007; Beaucourt *et al*. 2004; Biben *et al*. 2005; Biben and Misbah 2003; Fischer *et al*. 1978; Haas *et al*. 1997; Kantsler and Steinberg 2005, 2006; Mader *et al*. 2006; Rioual *et al*. 2004; Tamba *et al*. 2011).

The quest for understanding the fundamental interactions between the vesicle dynamics and viscous stresses in the surrounding medium motivated several early studies on the underlying transport phenomena in steady linear shear flows, both from theoretical (Beaucourt *et al*. 2004; Keller and Skalak 1982; Mader *et al*. 2006; Misbah 2006; Noguchi and Gompper 2007) and experimental (Abkarian and Viallat 2005; Christopher *et al*. 2008; Haas *et al*. 1997; Kantsler and Steinberg 2006) perspectives. These studies could unviel three distinct types of vesicle motion (depending upon the magnitude of viscosity ratio between the inner and suspending fluid and the shear rate), namely, Tank treading (TT) (Beaucourt *et al*. 2004; Biben *et al*. 2005; Biben and Misbah 2003; Rioual *et al*. 2004), Tumbling(TB) and an intermediate stage between TB and TT, alternatively known as vacillating breathing (Misbah 2006). These studies were complemented by experiments (Coupier *et al*. 2012; Vitkova *et al*. 2004) and theories on vesicle dynamics in pressure-driven flows as well, having conceptual resemblance with the physiological pumping. These studies could bring out unique shape transitions of the vesicle (bullet, croissant, parachute, and slipper), which could be related to its initial deflation (deviation in shape from sphericity) and the domain-confinement (ratio of vesicle size to channel width) of the exterior fluidic media.

In a realistic physiological microenvironment, the cellular matter does not encounter an idealized uniformity in the channel sections as considered in many of the early studies but rather confronts inevitable variabilities in the flow geometry, commonly resulting from typical patho-physical scenarios such as stenosis and aneurysms. Under such scenarios, adenosine triphosphates (ATPs) are known to trigger a vasodilatory signal toward controlling blood pressure as a natural physiological adaptive mechanism. In addition, metabolic abnormalities are also believed to be linked with the reduced deformability of RBCs upon entering a constriction (Zeng *et al*. 2016). Further, fluidic constriction in human vasculature has an established influence on the clustering of biochemically active particles, including drug delivery agents or activated platelets, bearing far-reaching consequences in forming blood clots (thrombus) in stenosed blood vessels (Bächer *et al*. 2017). Furthermore, constriction-induced elongation of cells is also linked with the mechanical transduction of biological signals, alternatively known as mechanotransduction, in choked flow passages (Mancuso and Ristenpart 2018; Wan *et al*. 2008). From a fundamental fluid dynamic perspective, all these scenarios inevitably feature abrupt constrictions in the flow passage that lead to the rapid convergence of the flow streamlines and thus result in drastic variations in the hydrodynamic shear stress in the exterior medium. The mechanism in which the encapsulated matter adapts to this dynamic alteration in the fluidic microenvironment depends critically on the viscosity contrast between the interior and the exterior fluids, the membrane fluidity, and the initial state from which it is released. While it renders unfeasible to study detailed hydrodynamics of the observed shape transition by probing human vasculature in-vivo, essential mechanistic insights of the same may be obtained via in-vitro bio-engineered microsystems on which explicit handles of flow control render to be feasible towards deciphering the input-output mapping unlike what could be done on living beings with challenged invasive procedures.

The advent of various microfabrication techniques over the recent years intensified the research endeavours towards in-vitro studies on the morpho-dynamics of deformable encapsulations in micro-environments having a high level of similitudes with the physiological systems under the intended design controls. Several of these studies focused on studying cellular dynamics in constricted microchannels. Particular examples include real-time deformability cytometry for a continuous mechanical characterization of biological cells via illumination and imaging (Otto *et al*. 2015), the dynamics of shear-induced ATP release from human RBCs (Wan *et al*. 2008), among others. These early studies were subsequently extended to characterize the mechanical behaviour of RBCs (Zeng and Ristenpart 2014) in pressure-driven flow, including their classical stretching behaviour (Mancuso and Ristenpart 2017).

Reported research reveals that while the dynamics of deformable encapsulations in a constricted passage have been examined extensively, the resulting inferences drawn appear to be mostly subjective and qualitative in nature. In other words, there needs to be more quantitative knowledge of their dynamic transition while passing across a constriction. This deficit stems from the complexities in providing precise experimental and theoretical evidence under controlled kinematic conditions, depicting the quantitative influences of the relevant geometrical and physical parameters on the dynamics of deformable encapsulations in a rapidly-converging fluidic environment.

Here, we capture quantitatively the unique role of the membrane bending rigidity and initial shape-deflation of a lipid vesicle as it transits along a geometrical constriction from stretching to tumbling motion with intermediate rolling under an applied pressure gradient. Our experimental and theoretical studies converge to a regime map that provides precise quantitative insight into these dynamic transitions in terms of the relevant normalized physical parameters. Furthermore, our study illustrates how that regime map may be utilized to estimate the membrane bending modulus of a vesicle by identifying the initial deflation and the observed dynamics as an alternative to established albeit complex rheometric routes. Our findings also bring out the effect of the initial transverse positioning of the vesicle, as well as the viscosity contrast between the inner and the outer fluid, on its extent of deformation. This quantitative depiction holds the potential of providing mechanistic insights into cellular aging (where the cytoplasmic viscosity increases with respect to normal healthy conditions) on its dynamic responses, for which the current state-of-the-art understanding is primarily empirical.

## 2. The physical problem: materials and methods

We consider a neutrally buoyant vesicle, filled with Newtonian fluid (density *ρ*_1_, viscosity *μ*_1_), suspended in another Newtonian fluid medium (density *ρ*_2_, viscosity *μ*_2_), and subjected to a parabolic flow profile with centerline velocity *u*_*c*_ in a converging microchannel, as shown in **Figure 1**. Both fluids are incompressible. A typical experiment starts with releasing an initially elliptical vesicle with unstretched major axis length *L*_*0*_, from an off-centerline position (nondimensional notation is eccentricity: *e* = 2 *y W*_*d*_) at the wider section (width, *W*_*d*_) of the channel and observing its dynamics as it migrates through the tapered section (*QR*) before the constricted channel (width, *W*_*c*_) as demarcated by the blue dotted line in Fig. 1. The constriction ratio of the channel is defined as *α* = (*W*_*d*_ /*W*_*c*_). The inlet (*PV*), outlet (*ST*), and wall (*PQRSTUV*) boundaries are demarcated as *S*_*i*_, *S*_*o*,_ and *B* respectively (see Fig. 1). The orientation angle (*θ*_*i*_) of the vesicle is defined as the angle between its major axis and the direction of flow. The Cartesian coordinate axis is located at the channel’s central axis. To gain insight into the quantitative variation in stretching, we define a parameter *λ*_*max*_ as the ratio of the maximum major axis length to the initial unstretched length of the major axis assuming a nearly ellipsoidal vesicle shape.

**Figure 1.**
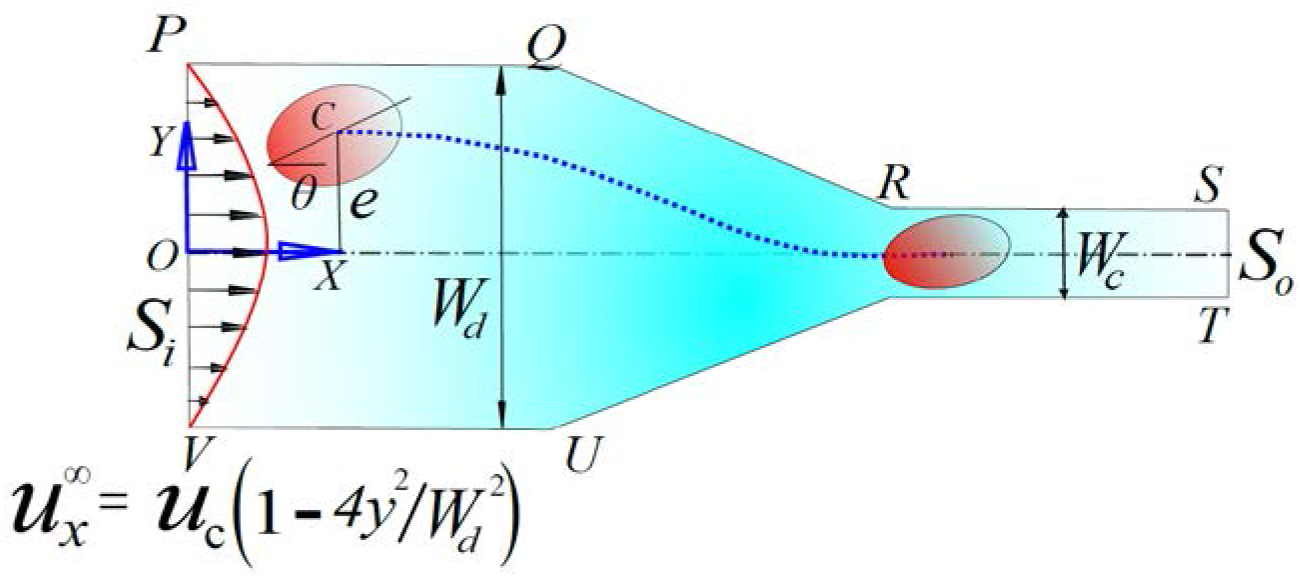
Schematic illustration of the flow domain depicting an initially elliptic vesicle, placed at an off-center position with eccentrically e, migrating through a constricted channel in plane Poiseuille flow at the inlet (*S*_*i*_). Typical parameters characterizing the vesicle are *L*_*0*_: the initial resting length; and *θ:* the orientation angle.

### 2.1 Numerical modeling

We use a combination of 2D and 3D numerical simulations to understand the different dynamic regimes (and their inter-transitions) of the vesicle morpho-dynamics, with an objective of confirming whether the essential physics can be captured using a 2D model that would otherwise avoid expensive computations.

#### 2.1.1 The 3D model

We deploy the Projective dynamics framework (Bouaziz *et al*. 2014), which is based on a Hamiltonian interpretation of the Newton’s laws of motion (Liu *et al*. 2017), to model the membrane of the lipid vesicle as a surface embedded in the 3-D space. In this method, the position of the interfacial marker points (obtained by triangulation of the surface) is advanced by minimizing the total energy of the vesicle, which is a combined consequence of the external forces on the membrane due to interaction with the carrier fluid and a its internal energies which arise from constraints enforced to represent the membrane properties of the vesicle. For a detailed discussion of the governing equations and the expressions for the internal energies, one may refer to Kotsalos et al. (Kotsalos *et al*. 2019).

The fluid flow is simulated using the Lattice Boltzmann method (LBM), where the evolution of the macroscopic flow variables such as velocity and pressure is modeled using the discrete velocity distribution function *f*_*i*_ (**x**, *t*) which evolves according to a set of collision and streaming rules that depend on the collision model and the lattice structure. Here, we employ a D3Q19 lattice structure where each lattice point is connected with 18 other lattice points lying inside the fluid domain. The Bhatnagar-Gross-Krook approach (Bhatnagar *et al*. 1954) is used for the collision operation, which updates the value *f*_*i*_ (**x**, *t*) towards an equilibrium value *f*_*i,eq*_ (**x**, *t*) according to the following equation

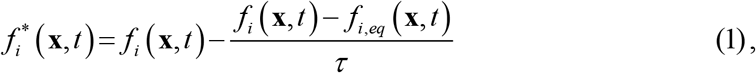

Where 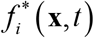 denotes the post-collision value of the velocity distribution function at a given lattice point, *τ* is the relaxation time parameter which is a function of the kinematic viscosity of the fluid and sonic speed in the medium. The streaming operation can be mathematically represented as,

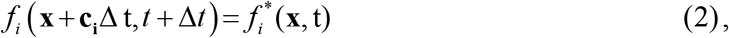

where **c**_**i**_ denotes the velocity vectors connecting two lattice points. The value of *f*_*i,eq*_ (**x**, *t*) at a lattice point is computed based on the value of the macroscopic fluid velocity and density at that point. A detailed discussion on the overall simulation approach can be found in the text of Kruger et al. (Krüger *et al*. 2017).

The external forces on the vesicle due to the fluid flow are lumped on the interfacial marker points. Values of the forces on these points are evaluated by interpolating the stress tensor of the fluid at the neighboring lattice points. The vesicle membrane is represented as an immersed interface; whose mesh is independent of the lattice structure of the fluid. The presence of the vesicle is communicated to the surrounding fluid using a force field, which ensures that the no-slip and no-penetration conditions are satisfied on the interface of the vesicle. This method belongs to a broad class of methods known as the Immersed Boundary Method’ which was developed originally by Peskin (Peskin 2002). Here, we apply this method using the Multi-Direct Forcing approach (Ota *et al*. 2012). The no-slip and no-penetration boundary conditions on the walls of the microchannel are enforced using the bounce-back method, where the collision step is replaced by a reflection step such that the fluid-particle populations hitting the bounce-back nodes are reflected along the opposite direction. This leads to an impermeable no-slip wall located exactly halfway between the bounce back node and the adjacent lattice point in the fluid. At the inlet of the microchannel, we enforce the fully developed velocity profile for Stokes flow, and at the outlet, a mass continuity is enforced.

In all the simulations, we set the dimensionless lattice spacing *dx* to be 0.5, the normalized kinematic viscosity of the carrier fluid *ν* to be equal to 1, and the relaxation parameter τ to be equal to 2. The size of the time step is obtained from the following diffusive scaling law:

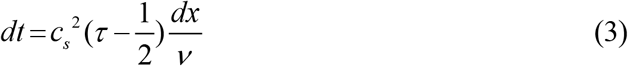

where *c*_*s*_ denotes the sonic speed. The stability of the LBM is ensured as τ > 0.5 (Krüger *et al*. 2017) and the results are found to be independent of the values of τ lying between 1 and 2. The material parameters for the vescicle are so chosen such that they fit the best with our experimental data for stretching along the axial coordinate (*λ* vs. *x*). The mapping between the simulation parameters and their corresponding physical values is mentioned in Table A1 of Appendix A. All the 3D LBM simulations are performed using the open-source PALABOS package (Latt *et al*. 2020).

#### 2.1.2 The 2D model

In an effort to explore computational economy, we employ the Boundary Element Method (Pozrikidis 2002) for our 2D simulations, which numerically evaluates the following expression for the velocity (*u*_*j*_) at a point (*x*_*0*_) lying on a boundary of the fluid domain:

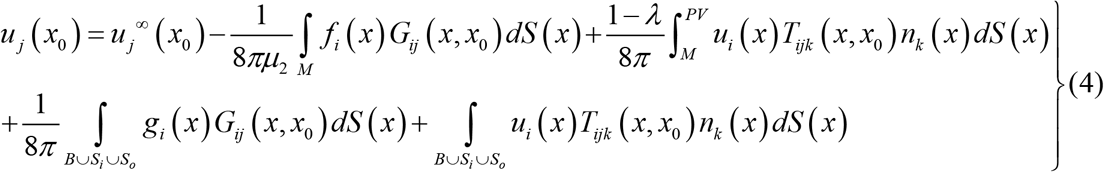

Here, *u*_*j*_^*∞*^ denotes the imposed velocity, *λ* is the viscosity ratio (*μ*_*1*_*/μ*_*2*_), *n*_*k*_ denotes the unit normal vector, *M* denotes the vesicle membrane, *S*_*i*_ and *S*_*o*_ denote the inlet and outlet of the channel respectively, *B* denotes the channel boundary and *G*_*ij*_ and *T*_*ijk*_ are the free-space Green’s function and corresponding stress tensor for Stokes’ equation (Pozrikidis 1992). The key term in equation (4), which accounts for the complex elastodynamic response of the membrane, is the vector *f*_*i*_(x) which denotes the jump in hydrodynamic traction at the vesicle membrane, and can be expressed as follows (Pozrikidis 2010):

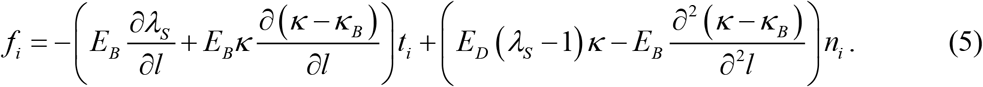

where *E*_*D*_ and *E*_*B*_ denote the dilatational and bending moduli of the membrane, *λ*_*s*_ denotes the local stretching of the membrane, *κ* denotes the local membrane curvature, and *κ _B_* is the membrane curvature at equilibrium. The arc length parameter is denoted using *l*, and the tangential and normal vectors are denoted by *t*_*i*_ and *n*_*i*_ respectively. The last two terms appearing on the right-hand side (RHS) of Eq. (4) denote the contributions to the interfacial velocity due to channel confinement. Since the velocity satisfies no-slip and no penetration boundary conditions at the walls, so the last term is identically zero on the channel walls. Further, since the channel inlet and outlet (*S*_*i*_, *S*_*o*_) are distanced sufficiently far from the vesicle interface, the double layer potential becomes vanishingly small as compared to the other terms (Leyrat-Maurin and Barthes-Biesel 1994). Therefore, the last term can be completely dropped off from the RHS of Eq. (4).

The traction exerted by the channel walls on the adjacent fluid is denoted by *g*_*i*_(x), which is unknown a priori. The channel traction (*g*_*i*_) is obtained by solving the boundary integral equation (4), on letting *x*_*0*_ be on the boundary (*B*U*S*_*i*_U*S*_*o*_). A similar approach was previously adopted for studying the dynamics of capsules (Leyrat-Maurin and Barthes-Biesel 1994) and droplets (Martinez and Udell 1990). The numerical algorithm initiates by assuming an initial shape and zero interfacial velocity at time *t*= 0. We then obtain the interfacial traction from Eq. (5). Based on the same, we solve Eq. (4) for the channel traction by letting *x*_*0*_ beto be located on the domain boundary. We next obtain the interfacial velocity from Eq. (4) by letting *x*_*0*_ to lie on the vesicle membrane. Subsequently, we advance the interfacial maker points using an explicit Runge-Kutta scheme. We then proceed back to the interfacial traction calculation using Eq. (5) and repeat the following procedure till the entire domain is traversed. We perform the entire computations using MATLAB, starting from the “*rbc_2d*” code available from the open-source BEMLIB package (Pozrikidis 2002).

### 2.2. Experimental procedure

#### 2.1.1. Vesicle synthesis

We synthesize the giant lipid vesicles (GUVs) in-house, following the standard electroformation (Angelova *et al*. 2007; Dimitrov and Angelova 1988) protocol at a room temperature of 27^0^ C. First, a lipid stock solution is prepared by adding 2.5 mg/ml of DOPC (Sigma-Aldrich) phospholipids to the chloroform-methanol solution (2:1 v/v). Then the stock solution is spread uniformly over an electrically conductive Indium-tin-oxide (ITO) coated glass (resistivity < 100 Ohm/sq. cm.; Sigma Aldrich) slide using a spin coater. 400 mM sucrose (Sigma-Aldrich) aqueous solution is used as the inner phase of vesicles. After drying of the stock solution under vacuum for 6-7 hours, the inner liquid is electro-swelled for 2 hours at a peak-to-peak voltage of 2.5-2.6 V_pp_, and a frequency of 10 Hz using an AC arbitrary waveform generator (Agilent, model no: 33250A) inside a closed chamber consisting of the ITO-glass electrodes separated by a non-conductive PDMS spacer (acting as a reservoir for the sucrose solution). This results in the formation of giant unilamellar vesicles of sizes ranging from 8 to 40 μm in diameter (Refer to the Supporting Information Figure S4. depicting vesicle size distribution). The vesicle solution is then diluted in slightly hypertonic glucose (Sigma-Aldrich) aqueous solution (435 mM concentration) to deflate the same by creating osmotic pressure difference and for obtaining the desirable contrast in refractive-index for visualization under phase-contrast mode.

An intended viscosity contrast between inner and outer fluid (different from unity) is achieved by adding Dextran (500 kDa, Hi-media) at 1%, 2%, and 3 % wt./ wt. to the inner sucrose solution. Please refer to Figure 2 depicting the alteration in shear viscosity of the inner solution of the vesicle as a function of % wt. of Dextran. The vesicle solution is then centrifuged (2 to 3 times) at a very gentle speed (15g-40g) for 30-35 minutes to sediment the vesicles at the bottom of the tube due to the action of centrifugal force. Centrifugation is important to wash out the outer medium and replace it with glucose aqueous solution of the same osmolarity (Kantsler and Steinberg 2006). During this process, the vesicles are handled carefully to avoid membrane rupture.

**Figure 2.**
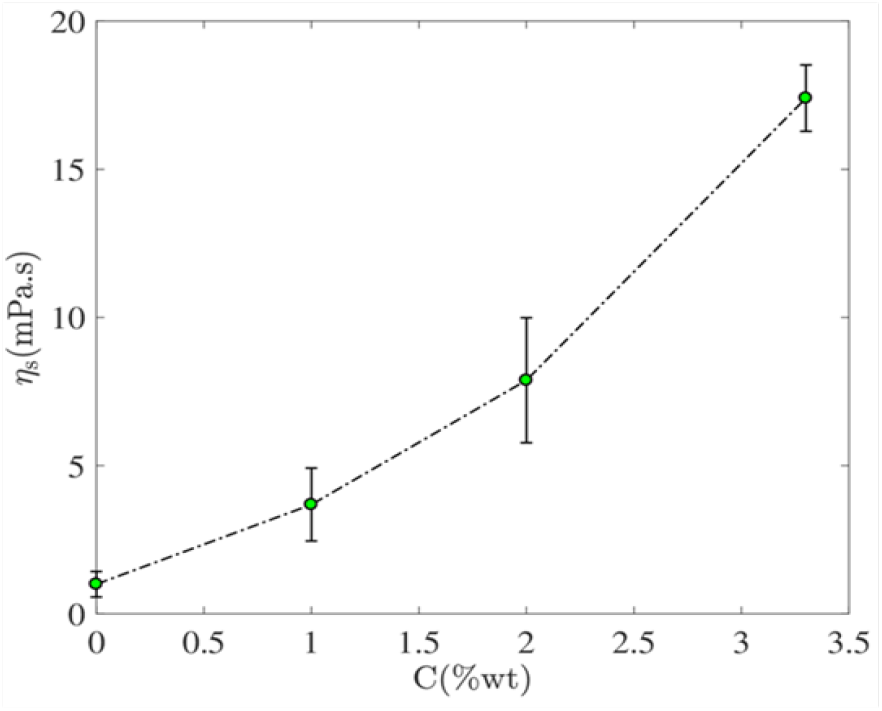
Variation in shear viscosity of the vesicle inner solution (containing Dextran) as a function of Dextran concentrations at 25^0^ C obtained from the rheometry result for the experimentation related to the viscosity contrast study.

The shear viscosities of the inner and the suspending phase liquids are measured using a stress-imposed rheometer (Anton-Paar, MCR 302; TA Instruments) running in the cone and plate configuration. All the measurements are made at a constant and controlled temperature of 25 ± 0.6 °C with the help of a Peltier temperature controller. The properties of the reagents used as inner and suspending phase fluids are enlisted in Table 1 below.

**Table 1.** Properties of the working fluids used in lipid-vesicle synthesis

| components | Chemicals used | Viscosity | Density |
| --- | --- | --- | --- |
|  |  | (mPas) | (Kg/m<sup>3</sup>) |
| *Inner-fluid* | Case-1 400 mM sucrose-Aq. solution | 1.06 | 984 |
|  | Case-2 400 mM sucrose-Aq. solution+1% w/w Dextran | 3.685 | 980 |
|  | Case-3 400 mM sucrose-Aq. solution+2% w/w Dextran | 7.88 | 987 |
|  | Case-4 400 mM sucrose-Aq. solution+3.3% w/w Dextran | 17.4 | 989 |
| *Outer-fluid* | 435 mM glucose-Aq. solution | 0.989 | 980 |

To alter membrane bending rigidity modulus, Glutaraldehyde (Hi-media) aqueous solution is added to the electroformed vesicle solution in a range of 0.5% to 10 % volume/volume. (Forsyth *et al*. 2010a). An inverse mapping technique is employed to estimate the membrane bending modulus from the maximum stretching versus the normalized bending modulus curve by superimposing the simulation and experimental results and converging them via an error minimization technique. Refer to the Supporting Information, Figure S3, and Table S1, for more details.

#### 2.2.2. Microfluidic experiments and flow visualization

The microfluidic channels are fabricated using PDMS as the base material, mixed with a cross-linker at a ratio of 10:1, following standard soft lithography protocol from the master pattern obtained via conventional photolithography technique (negative photoresist SU8 2050, Mirco-Chem). Post-curing, the microchannels are plasma bonded (Harrick plasma) to a glass coverslip to facilitate the experimental study and flow visualization. Experiments on the microfluidic test bench are executed at a controlled temperature of 25 ^0^C. The constriction ratio and flow rates are combinedly altered to impose precise variations in the extensional strain on the deforming encapsulations (refer to the experimental setup in **Figure 3**). Representative widths of the microchannel at the upstream of the convergent section (*W*_*d*_) and the constricted section (*W*_*c*_) are ∼ 100 μm and ∼ 80 μm, respectively. The height of the channel was maintained *h* ∼ 30 ± 3 μm. The convergent section has a typical length of 200 μm out of a total axial length of the test rig of about 3 cm. The distance from the inlet to the converging section is ∼1 cm, which is large enough as compared to the hydrodynamic entry length in the Stokes flow limit.

**Figure 3.**
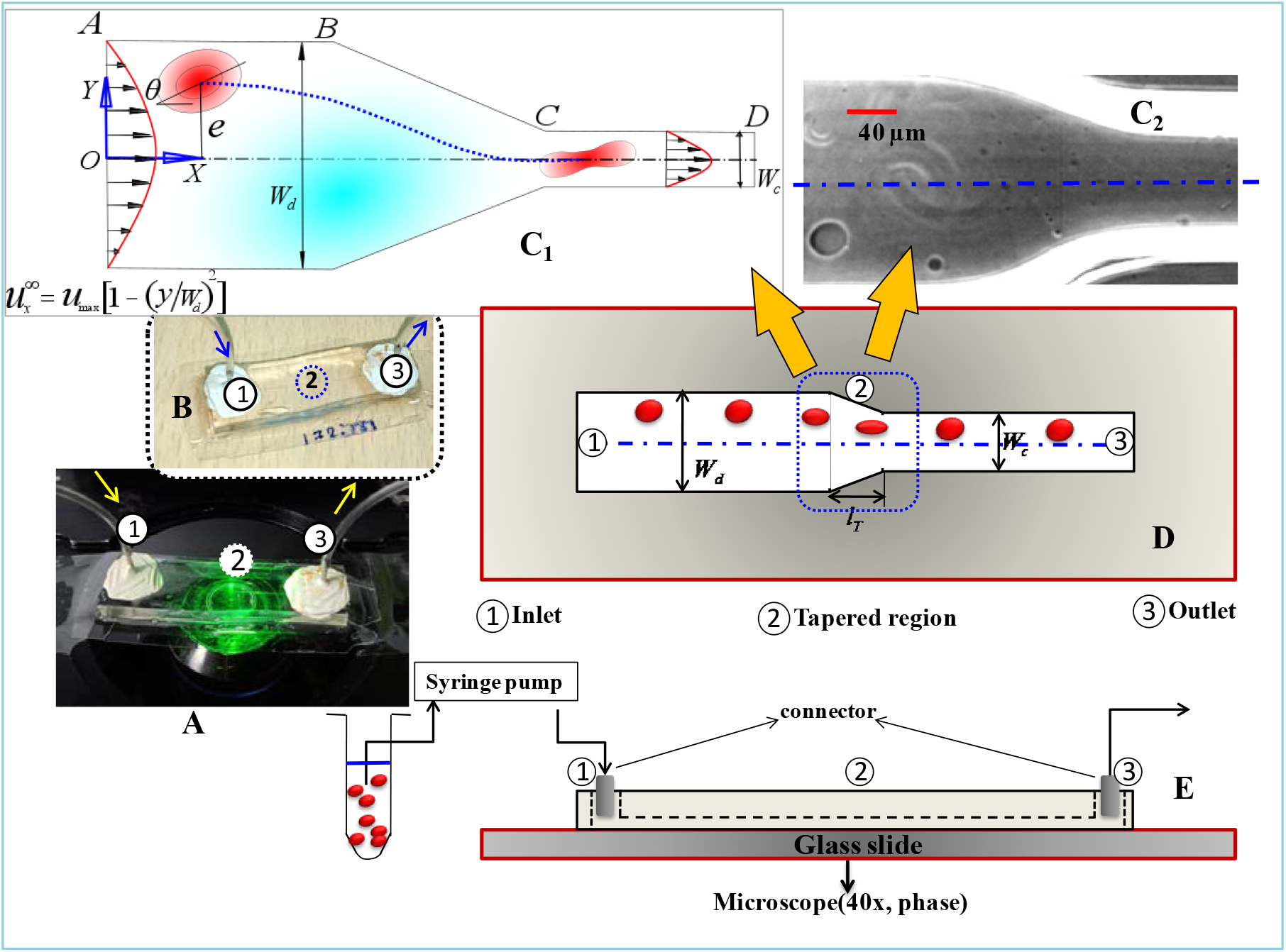
The physical system: (A), (B) Experimental setup having flow inlet (1), outlet (3), and the test section (2). The transport phenomena are observed under an inverted transmission-type microscope operated in phase-contrast mode. The direction of flow is shown using arrows. Panel C_1_ shows a schematic of the flow domain. W_d_ and W_c_ denote the width of the diverging section (AB) and constricted section (CD), respectively. BC denotes the tapered region. The coordinate axis is located at the channel centerline. The normalized distance of the observed cell centroid from the channel centerline is denoted by eccentricity *(e). θ* denotes the initial inclination angle of the major axis of the cellular outline with the flow direction. Panel C_2_ shows the experimental test section (denoted by 2 in Fig. 3A, 3B). Panels D and E denote the schematic of the microfluidic channel and the experimental setup. The Blue dotted line in panel D denotes the test section.

To begin the experiment (see **Figure 3**), the vesicle suspension is first loaded in an air-tight Hamilton glass syringe, and any existing gas bubbles are purged out. Next, the syringe is connected to a syringe pump (model number PHD 2000, Harvard Apparatus) operating in the infuse mode to inject the sample at the microfluidic test section at a range of controlled flow rate (*Q*) ∼ 30 to 180 μL/hour [*Re* ∼ *O* (10^−2^-10^−1^)]. Polyethylene tubings (0.58 mm inner diameter) are used to connect the syringe to the inlet of the microfluidic device to complete the fluidic circuitry. The flow rates are varied over the range *Q* =30 μL/hour to *Q*=180 μL/hour respectively. The respective shear rates, 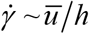 (where *h* is the channel height, and *u* is the cross-sectionally averaged flow velocity) are ∼10 s^-1^; and ∼100 s^-1^. The vesicle dynamics is visualized with an inverted microscope (Olympus IX71) operated in phase-contrast mode, (40X magnification) coupled with a high-speed camera (Phantom-v7) to probe the same under altered dynamic conditions. The high-speed camera is set to capture time sequence images at a frequency of 6000-10200 Hz. The obtained time-series image sequences are subsequently interpreted using a MATLAB GUI (Basu 2013) to obtain the contour, centroids, trajectory, velocity, and orientation angle of the moving object.

## 3. Results and Discussion

The observed vesicle dynamics can be mapped to one among three regimes, namely “stretching”, “rolling” and “tumbling”. In the stretching regime, the vesicle undergoes an elongation about its major axis as it passes through the constriction and thereafter relaxes to an equilibrium shape. On the other hand, the tumbling regime is characterized by a rotation of the vesicle about an axis perpendicular to the plane of flow and is characterized by negligible shape elongation. In the “rolling” regime, the vesicle exhibits characteristics of both stretching and tumbling as it is associated with a simultaneous shape elongation and rotation about its major axis. While the above shapes were qualitatively observed in the past, we aim here to put forward their quantitative depiction as mapped with the relevant physical properties. Towards this, we identify two important non-dimensional parameters responsible for these shape transitions, namely: **[a]** reduced area (*τ* _2*d*_) or, reduced volume(*ν*) applicable for 2D or, 3D analysis expressed as *τ* _2*d*_ = *A/π* (*p*/2*π*)^2^ or, *ν* = [3*V*/ 4*π* (*A*_*s*_/ 4*π*)^3/2^] respectively, which is the ratio of actual surface area (*A*)/ volume (*V*) of the vesicle to the area of a circle/ sphere having the same perimeter (*p*)/surface area(*A*_*s*_), and **[b]** normalized bending-rigidity-modulus 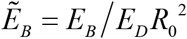 where *E*_*B*_ is the modulus of membrane bending rigidity, *E*_*D*_ is the dilatation modulus of the vesicle membrane and *R*_0_ is the effective radius of the vesicle, which is defined to be the radius of a circle having an equivalent surface area and can be expressed as 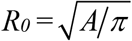 (for 2D) or, 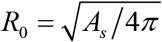 (for 3D). The parametric dependence of *τ*_*2d*_ and 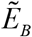 on the shape dynamics is elucidated with the help of a regime diagram. We observe that the transition from stretching to tumbling can be identified as belonging to a well-known family of dynamical transitions known as saddle-node bifurcation. Another important dimensionless parameter is the confinement ratio, *C*_*n*_ = 2*R*_*0*_/*W*_*c*,_ defined as the ratio of the effective radius *R*_*0*_ of the vesicle to the half-width of the constricted channel. We also quantify the effect of the ratio of cytoplasm (inner) to extracellular (outer) fluid viscosity: *η*_*r*_= *η*_*in*_*/η*_*out*_ on the extent of stretching. As quantified by stretch ratio, *λ* = *L/L*_*0*_, where *L* is the instantaneous major axis length and *L*_0_ is the initial major axis length of an equivalent ellipsoid-shaped geometric entity. The maximum stretch ratio (*λ*_*max*_), is accordingly obtained as the ratio of the maximum instantaneous distance (*L*_*max*_) between a point on the vesicle contour and its centroid to its initial/unstretched distance (*L*_0_).

Our computational studies are aimed to mimic the experiments with a flair of property variations having established similitude with the movement of RBCs. For the same, we consider the reduced area to vary between 0.6 and 1 (Abkarian *et al*. 2007; Dupire *et al*. 2012), and normalized bending modulus 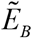 to vary between *O*(10^−6^) – *O*(10^−2^). These normalized parameters physically correspond to established membrane properties of vesicles and RBCs : *E*_*B*_ ∼ 3×10^−19^ J ; *E*_*D*_ ∼ 10^−7^ −10^−1^ N m ; *R*_*0*_ ∼3-40 μm (Pozrikidis 2010). The computational values of *v* range between 0.85 to 0.99, conforming to our physical experiments. All the results are obtained in the creeping flow regime conforming to low Reynolds number (*Re* = *ρ*_*out*_*V*_*av*_*W*_*c /*_*η*_*out*_) hydrodynamics, where the value of *Re* ∼ O (10^−2^ - 10^−1^) is calculated based on the extracellular fluid property.

### 3.1 Stretching (S), rolling (R) and tumbling (T) dynamics prior to the constriction

**Figure 4(a-c)** illustrates different morpho-dynamic features as the lipid vesicles pass through the tapered region before the constriction (refer to the schematic of Fig. 4d). While describing these motions, the vesicles are considered without any viscosity contrast (*η*_*r*_ ∼ 1), to bring out exclusive influences of the bending rigidity of the membrane. In the stretching motion (Fig. 4a), the vesicles experience geometric constriction induced elongational strain due to the reduction in the cross-sectional area resulting in a smooth elongation in their shape in the direction of flow, quantified by stretch-ratio (*λ*). This reaches a maximum at the end of the tapered region (corresponding to the maximum extensional rate) and then gradually reduces to a steady value as it reaches an equilibrium shape inside the constricted channel. The insets in Fig. 4a delineate the vesicle shape as a function of axial distance for a representative vesicle of reduced volume ∼ 0.99 corresponding to a shear rate of ∼ 210 ± 3 s^-1^ at the constricted section. The magnitude of the shear rate at the constricted section is maintained nearly the same as in the case of stretching motion by adjusting the flow velocity and the transverse position of the vesicle. The effective radius, for the representative vesicle, is *R*_*0*_ ∼ 11.12 *μm*. In the absence of any viscosity contrast between the inner and the outer fluids, it is the confluence of membrane bending rigidity modulus, 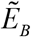 and initial shape deflation(*τ* _2*d*_) that controls the dynamic transition (see the regime plot, Figure 5a). Figure 3c represents the tumbling motion of a vesicle (*ν* = 0.965 ; *R*_0_ = 11.25 *μm*), akin to rigid-body rotation (see Fig. 4e) represented by the variation in its orientation angle (*θ*) along the axial position, exhibiting characteristic spatio-temporal discontinuity. The rolling motion features a continuous periodic variation of *θ* versus *x/R*_*0*_ *(ν* = 0.915 ;*τ* _2*D*_ = 0.92 ; *R*_0_ = 11*μm)* (see Fig. 4b), showing the combined characteristics of stretching and tumbling, akin to saddle-node bifurcations where the fixed points are either created or destroyed along with a parametric sweep.

**Figure 4.**
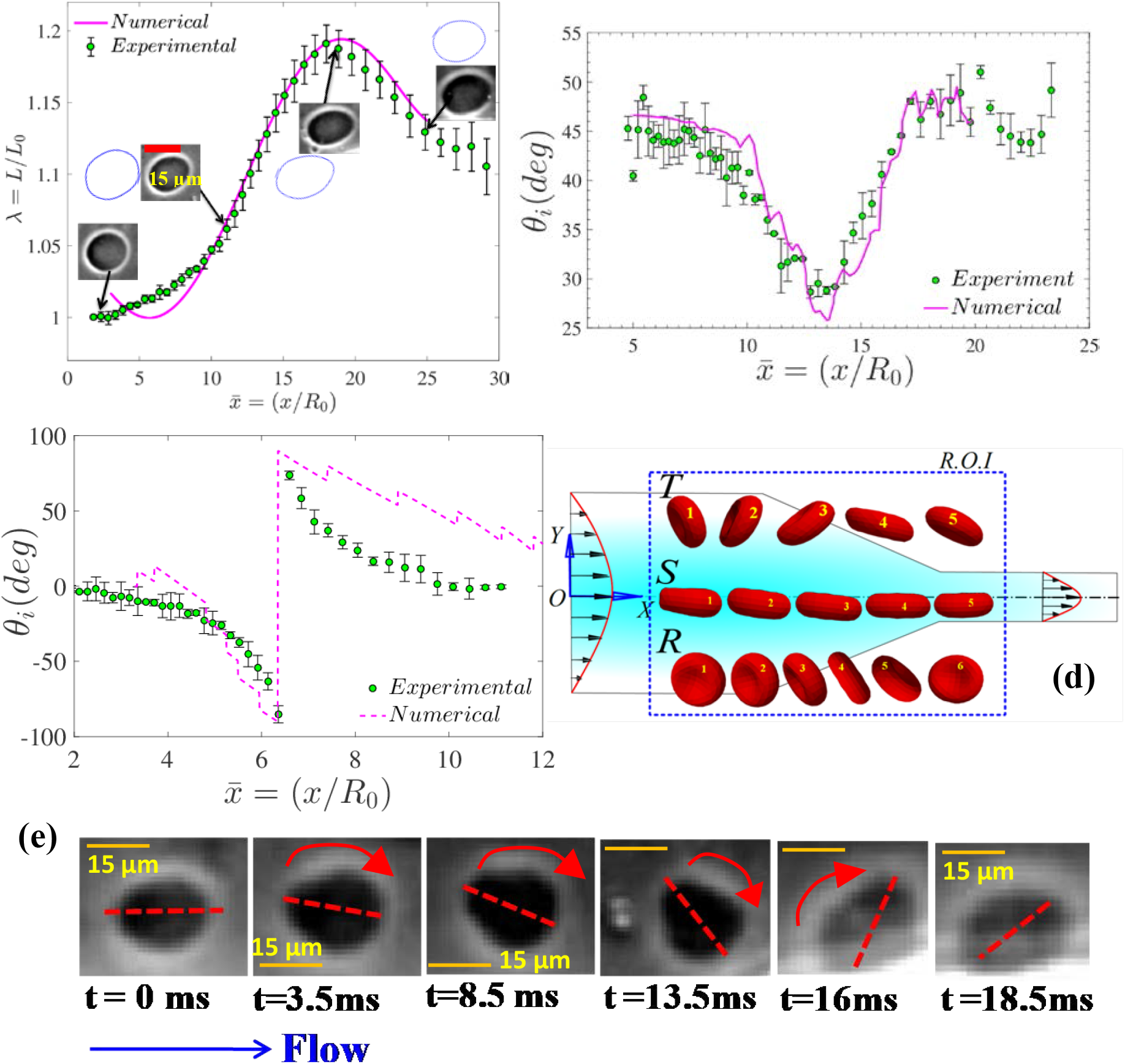
Three different vesicle dynamics were observed prior to the constriction: **(a)** stretching dynamics demarcated by stretch-ratio (λ) as a function of normalized axial displacement obtained from the controlled experiment (for*ν* = 0.99) and 2D numerical simulation (*τ* _2 *D*_ = 0.99). The insets represent the vesicle shapes obtained from the experiment and 2D simulations side by side at different axial locations. **(b)-(c)** Evolution of the vesicle orientation angle (in degree) as a function of normalized axial displacement obtained from experiment and numerical simulations for vesicles undergoing rolling (*ν* = 0.915 ;*τ* _2*D*_ = 0.92), and tumbling (*ν* = 0.965 ;*τ* _2*D*_ = 0.96) motions respectively. Open symbols denote the experimental results while solid/dotted lines represent the results obtained from numerical simulations. The confinement ratio is maintained ∼ 0.25-0.27 for all three cases. **(d)** Schematic representing the three different motions experienced by vesicles while passing through the converging section prior to the constricted channel. **(e)** Time-lapse images showing the temporal evolution of the orientation angle of a vesicle undergoing tumbling motion obtained from experiments. The red arrow denotes the sense of rotation, where the axis of rotation is perpendicular to the plane of the paper. The scale bar is mentioned on each experimental view graph. Please refer to the Supporting information (SI), movies S1-S3 for more details.

**Figure 5.**
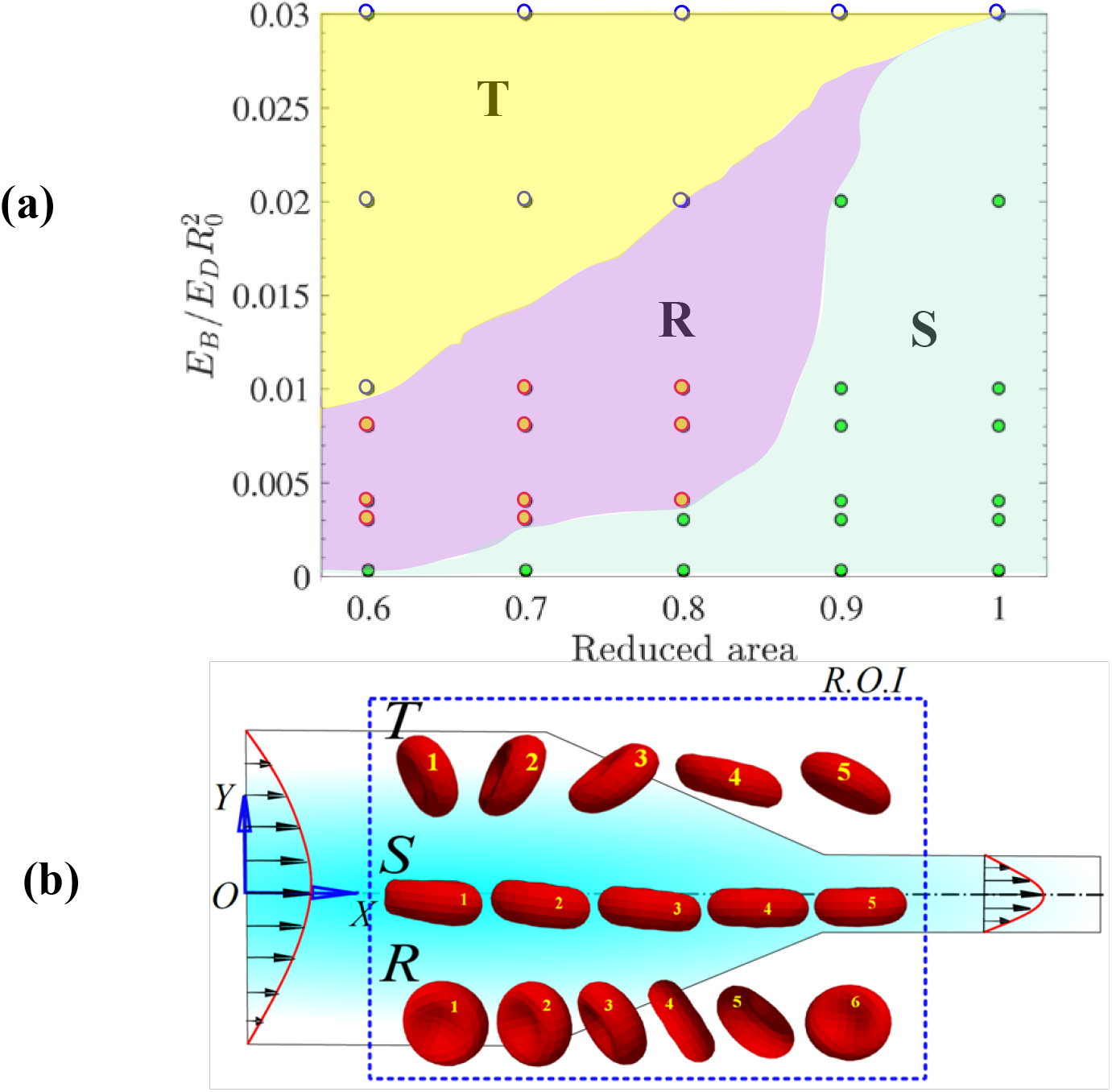
**(a)** Transition of motion features as a function of reduced area and normalized membrane bending-rigidity modulus. The S (stretching), R (rolling), and T (tumbling) regimes map quantitatively with the membrane rigidity. The same, along with the shape alterations over the stretching regime, enable quantitative prediction of the mechanical properties of the vesicle membrane without resorting to invasive probing **(b)** the schematic depicting observed dynamics over space and time, corresponding to the average shear rates at the channel constriction ∼ 208 ± 3 s^-1^. All the data points representing the regime map is obtained from 2D simulations and validated with controlled experiments.

On comparison with the numerical simulation results, it is evident from Figs. 4(a-c) that our 2D model is in good agreement with the experimental findings. These results, and sweeps over broad parametric spaces, evidence that the physics of shape evolution is reliably captured via 2D computations and in good agreement with the 3D model (see *Supplementary Information*, section 1 for more details). Therefore, to arrive at an optimal balance of physical consistency and computational economy, 2D numerical simulations may be adhered to.

### 3.2 Vesicle membrane stiffness mediated S-R-T transition in the microfluidic constriction

We further traverse the entire dynamic regime starting from a stretching regime, with controlled alterations in the vesicle membrane stiffness via an optimized experimental protocol. For example, in an effort to stiffen the vesicle membrane against bending, these are first incubated for 20 minutes at room temperature (∼26 °C) (Forsyth *et al*. 2010b) with 5% to 10% vol./vol. glutaraldehyde (Sigma-Aldrich) solution and then used immediately for experimentation. The addition of glutaraldehyde helps polymerizing the lipid membrane and thus altering its bending rigidity modulus to an extent that it can arrest the extent of stretching to a desired extent. The vesicles having controlled variations in the membrane stiffness, thus functionalized, accordingly exhibit different extents of stretching (refer to the SI, Figure S3).

**Fig. 5(a)** showcases the different dynamic regimes that are traversed in the process, for a representative scenario having no viscosity contrast between the inner and the outer fluid, subject to an average shear rate of about 209 ± 2 s^-1^ at the constricted region of the channel. The results clearly demonstrate the interplay between the membrane bending rigidity modulus, 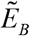 and its initial shape deflation towards transiting from one dynamic regime to another. From Fig. 5(a), it may be observed that for 0.6 <*τ* _2 *D*_ < 0.936, an increase in 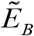 beyond a threshold limit results in a smooth transition from stretching to rolling to rumbling motion, for a given shear rate. For lower values of 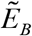, the membrane can restore to its original shape upon removal of external stress via elastic recovery (Skotheim and Secomb 2007). For sufficiently larger values of 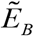, the elastic energy rise may endure tumbling as an energetically more favourable proposition as compared to the energy-expensive stretching. For the intermediate values of 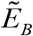, combined rotation and stretching is accompanied by a smooth variation in the orientation angle under rolling motion. Quantitatively, the resulting conformation alterations are manifested by a continuous change in the orientation angle *θ* made by the major axis of the vesicular encapsulation with the flow direction with reference to its normalized axial position. The corresponding numerical predictions agree well with the experimental trends (see Figure 4).

The observed shape transitions may be rationalized by introducing a parameter *ϕ*(*t*), denoting the instantaneous phase angle of an infinitesimal membrane element, so that its time rate of change, *dϕ*/ *dt*, may be linearly mapped with the frequency of the observed motion (Keller and Skalak 1982). Accordingly, the following inequalities hold: 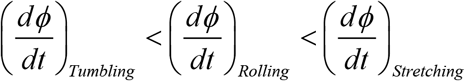. For a model encapsulation having an ellipsoidal boundary (Keller and Skalak 1982), one may get: 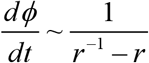, where *r* is the ratio of the minor axis to the major axis length of the ellipse. Physically, a decrease in *r* would implicate declining *dϕ*/ *dt*, resulting in a reduction in the frequency of equivalent tank-treading motion. A sharp increase in the tank-treading frequency may further be observed as the value of *r* approaches unity, leaving apart the singularity at *r* =1. Further, for a fixed magnitude of the bending modulus of the membrane, since *τ* _2*d*_ decreases with a decrease in *r* from pure geometric considerations, it follows that *dϕ*/ *dt* also decreases with a decrease in*τ* _2*d*_. From Figure 4(a-c), it may be observed that the orientation angle (*θ*) remains virtually constant over the stretching regime (refer to SI, *Fig. S1a*) but exhibitsa discontinuous yet periodic behavior during tumbling. Hence, both stretching and tumbling events are fixed point solutions in the (*ϕ* - *θ*) plane, and their basins of attraction may be demarcated exclusively by two parameters, namely, the normalized bending modulus and the degree of initial deflation (see Fig. 5a).

### 3.3 Effect of initial off-center position on the stretching dynamics in the constriction

**Figure 6a** delineates the effect of the initial transverse position of a vesicle, represented by its eccentricity (*e*), on its maximum stretching while keeping the other parameters fixed. The maximum stretch ratio (*λ*_*max*_) shows a nonlinear, monotonically increasing trend with an increase in eccentricity (*e*), as depicted in the figure.

**Figure 6.**
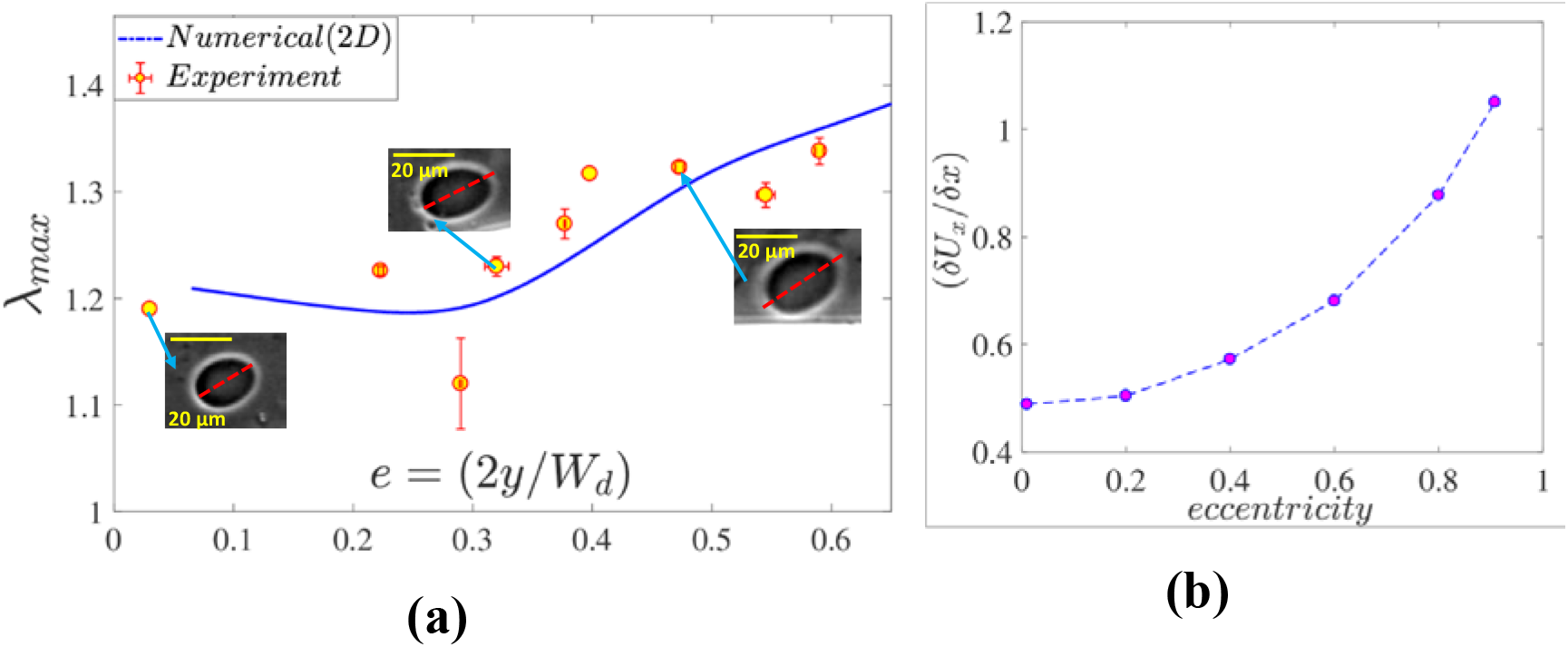
**(a)** Variation in maximum stretch ratio as a function of normalized initial transverse position from the channel centerline (eccentricity) for a vesicle undergoing stretching dynamics. Open symbols demarcate experimental results, while solid lines are obtained from numerical simulations. The insets in Fig. 5a denote the experimental view graphs corresponding to fixed eccentricity values. **(b)**Variation in axial velocity gradient/ extensional rate with the normalized lateral distance from channel centerline (eccentricity). The other important parameters are *ν* = 0.985; *R*_0_ = 13.5 *μm* ; *τ* _2*d*_ = 0.986.

An explanation of this trend follows from Figure 6(b), which represents the variation of normalized axial velocity gradient (*δU*_*x*_*/δx;* strain rate) as a function of the transverse position of the vesicle within the constricted channel. Further, this axial velocity gradient is proportional to the extensional stress acting on the vesicle membrane resulting in its elongation along the flow direction. From Fig. 6(b), we can conclude that the magnitude of extensional stress on the vesicle membrane increases with an increase in eccentricity(*e*), which clearly explains the trend of *λ*_*max*_ versus eccentricity as observed in Fig. 6(a). The corresponding results obtained from numerical simulations are in good agreement with the experiments as well.

### 3.4 Effect of viscosity contrast on stretching dynamics

**Figure 7a** depicts that for a constant shear rate (208 s^-1^ at the constricted region), the maximum stretch ratio (*λ*_*max*_) increases sharply with a marginal increase in the inner-to-outer fluid viscosity ratio *η*_*r*_, till an optimal value (*η*_*opt*_) ∼ 5.8 may be reached. Beyond that limit, *λ*_*max*_ attains an asymptotic saturation. The inset located at the right-hand side panel of Fig. 7(a) shows the variation in stretch ratio as a function of its axial position for two different viscosity contrasts (*η*_*r*_ =1 and 8). The experimental and simulation viewgraphs in Figure 7a represent the vesicle contours at the different axial positions while moving through the converging section of the microfluidic channel before the constriction. It is evident from Fig. 7(a) that the simulation results agree quite well with our experimental findings.

**Figure 7.**
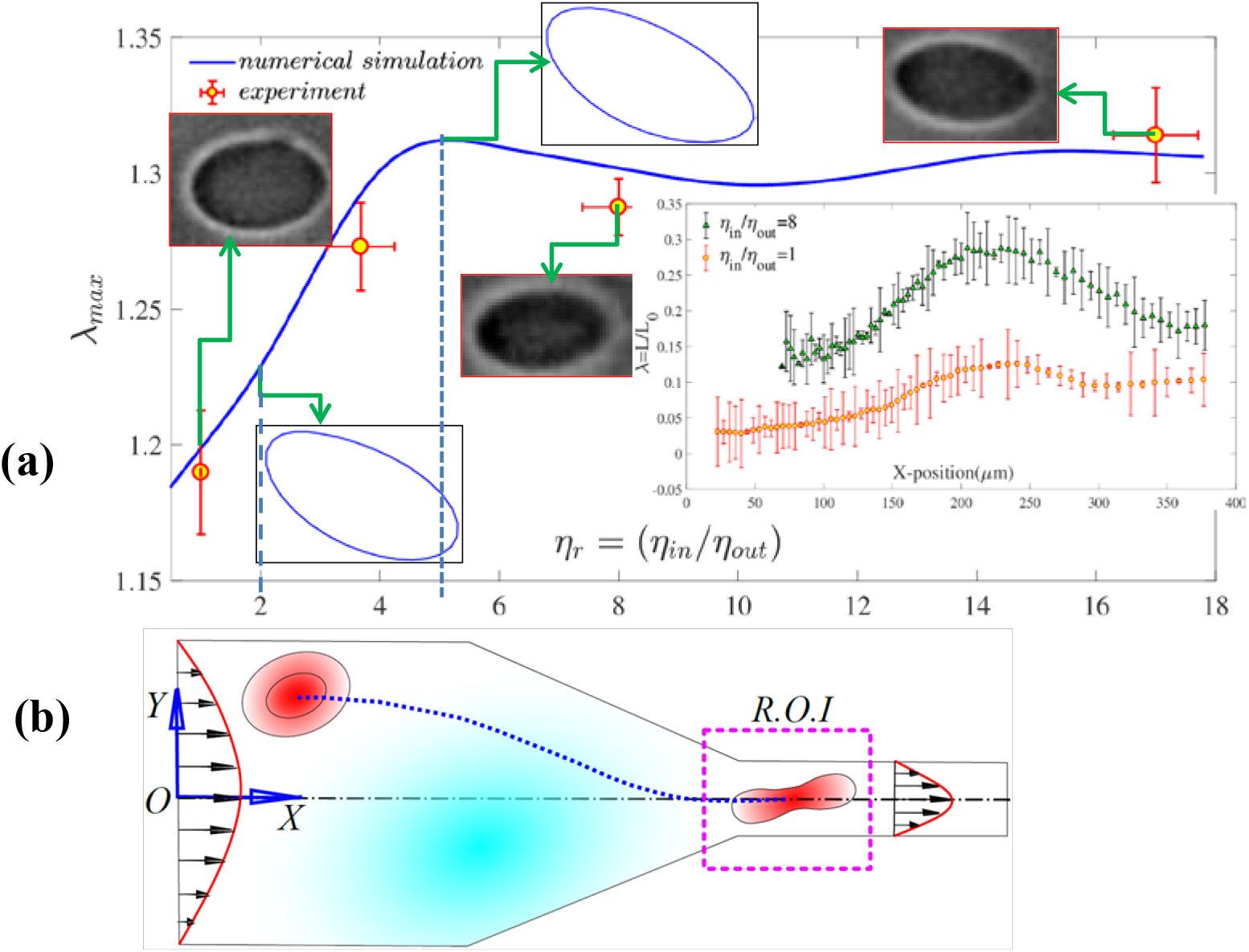
**(a)** Variation in the maximum stretch ratio (*λ*_*max*_) as a function of viscosity contrast (*η*_*r*_) between inner and outer fluids of the vesicle, as obtained experimentally (reduced volume ∼0.95, *C*_*n*_ ∼ 0.21) as well as from 2d-numerical simulations (reduced area∼0.94, *C*_*n*_ ∼ 0.2). The shear rate at the constriction is maintained ∼ 208 s^-1^. The initial off-center position of the vesicle is ∼ 0.67. Open symbols demarcate experimental results, while blue solid lines represent numerical simulation results. The inset located at the right-hand side of the main panel shows the variation in the stretching ratio (*λ*) as a function of its axial position for two different viscosity contrasts: *η*_*r*_=1, 8. The view-graphs in the main panel demarcate the contours of the stretched vesicle at its maximum limit for different viscosity ratios). **(b)** Schematic representation of the region of interest where the maximum stretching takes place.

The increase in *λ*_*max*_ with *η*_*r*_ (Figure 7a) can be explained in the light of excess viscous stress exerted by the inner fluid on the lipid bilayer membrane of the vesicle, leading to subsequent stretching, and deformation of the bilayer membrane. Since the viscous stresses scale linearly with the fluid viscosity, the same would intuitively facilitate stretching. However, on increasing *η*_*r*_ beyond *η*_*opt*_, the fluid circulation inside the vesicle gets attenuated to the extent that the same behaves effectively like a rigid encapsulation. For a quantitative mechanistic insight on the same, one may refer to Figure 8, which reveals that for a fixed position on the interface (corresponding to a fixed polar angle; *θ*), the magnitude of the traction force increases with the increase in viscosity ratio, realizing an enhanced stretching ratio with an increase in *η*_*r*_ till *η*_*opt*_ is reached.

**Figure 8.**
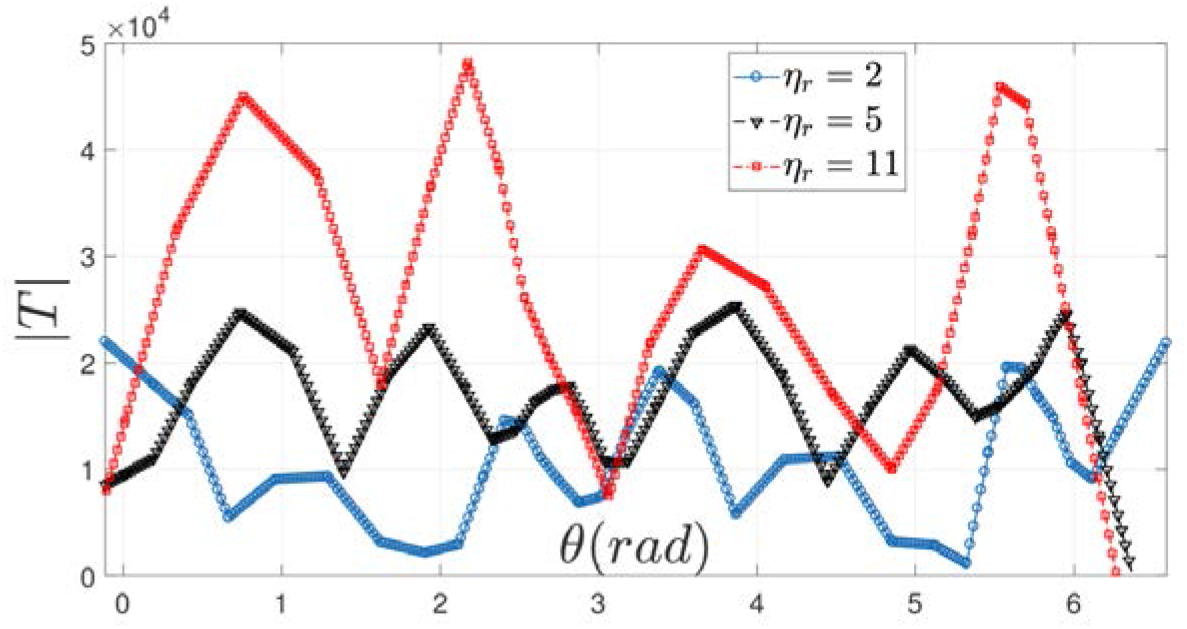
Variation in the magnitude of traction force (|***T***|) on the vesicular interface as a function of the polar angle (*θ*) for 3 different viscosity ratios, as obtained from numerical simulations. The reduced areas corresponding to *η*_*r*_ =2,5 and 11 are *τ* _2*d*_ ∼ 0.94. *C*_*n*_ for all three cases is ∼ 0.21.

The above results obtained via modulation of the viscosity ratio appear to imperative in explaining the altered dynamics of the RBCs in human vascular pathways with ageing, on the account of an increase in cytosolic viscosity (Ma *et al*. 2022) due to the increased concentration of haemoglobin molecules, to an extent up to 3 to 5 folds with reference to the average value of the same over the healthy life cycle of the RBC (∼5.5 mPa.s). Similar phenomenology may manifest in certain diseased conditions (for example, malaria) due to the polymerization of haemoglobin as a common pathological artefact in the infected RBCs. While these observables are reported as potential disease markers in medical literature, their fundamental mechanco-physical routes, which remained elusive thus far, may be potentially traversed by the approach outlined in this work.

## 5. Conclusions

We investigated the dynamics of singled-out, deformable lipid vesicles through a constricted microfluidic pathway. The results obtained from the numerical simulations were validated with in-house experiments deploying laboratory-synthesized lipid vesicles, showing good agreement. The particular focus was to quantify the mapping between the membrane properties of the vesicle and its initially deflated configuration towards the transition of motion from stretching to tumbling via rolling. The rolling motion combined the characteristics of both stretching and tumbling dynamics which could be precisely quantified by the pertinent morpho-dynamic parameters and their evolution with time. The stretching dynamics was shown to be further modulated by altering the initial position (eccentricity) of the vesicle with respect to the central axis of the channel. The consequent variations in the stretching ratio exhibited a nonlinear, monotonically increasing trend with eccentricity due to increasing strain rate. Finally, we examined the effect of viscosity contrast of the inner and outer fluid over the stretching regime. The maximum stretching ratio, in the process, was shown to increase by about five times, before attaining saturation. This could hallmark the transition of the flexible vesicle to a nearly rigid entity that inhibited further shape transitions.

In addition to providing quantitative mechanistic insights on the vesicle dynamics, we believe that our study could act as a prelude to explaining various cellular morphologies observed in human microvascular physiology. While qualitatively, many such observations on altered morphology of different cells, typically RBCs, were previously reported as patho-physical alterations on account of various diseases or aging, no effective quantitative rationalization on the same could be arrived at because of a lack of precise mapping between the geometrical and physical properties and the observed morphologies, rendering the state-of-the-art on the same to be primarily phenomenological and empirical rather than fundamental. In that perspective, our results offer natural promises of establishing a quantitative connection between the observed cellular features and their properties, hallmarking various aspects of health and disease. Proceeding further forward, with unprecedented recent advancements in high-speed and high-resolution imaging, photographs of the dynamic images of identified cells may be potentially utilized to predict the mechano-physical properties of the same under dynamic conditions as an alternative to traditional rheometry that may not depict the realistic straining conditions as a living cell traverses complex physiological pathways having inevitable undulations and constrictions.

## Acknowledegments

T.K. wants to thank Mr. G. Coupier (CNRS researcher at Liphy, France), Prof. M. Abkarian (Montpellier Structural-Biology-Center), Prof. C. Misbah (Director of research at CNRS, France) and Prof. V. Kantsler (University of Warwick, UK) for sharing their insightful suggestions via email. T.K wants to convey his sincere gratitude to Prof. Sovan Lal Das (IIT Palakkad) for his help on the lipid vesicle synthesis via electroformation method at the early phase of learning. T.K. acknowledges helpful discussions with Mr. Banuprasad T.N (Research Scholar, IIT Kharagpur) regarding experimental issues. D.P.P. is thankful to Dr. Christos Kotsalos for his help in the implementation of the PALABOS library. This work needed the supercomputing facility of IIT Kharagpur, established under the National Supercomputing Mission (NSM), Government of India, and was supported by the Centre for Development of Advanced Computing (CDAC), Pune.

## Funding Statement

S.C thankfully acknowledges the SERB, Department of Science and Technology, Government of India, for Sir J. C. Bose National Fellowship.

## Declaration of Interests

The authors declare no conflict of interest.

## Author Contributions

T.K., D.P.P, and S.C. designed the research problem. T.K. performed all the experiments and image analytics. D.P.P performed numerical simulations. T.K., D.P.P, and S.C. analyzed the data and wrote the paper.

## Data Availability Statement

All study data are included in the article and *supporting information (SI)*.

## Ethical Standards

The research meets all ethical guidelines, including adherence to the legal requirements of the study country.

## Appendix A

**Table A1.** Mapping between simulation parameters and their physical values for 3D numerical simulation.

| Parameters | Numerical value | Physical value |
| --- | --- | --- |
| Grid Spacing ($dx$) | 0.5 | 0.5 $\mu\text{m}$ |
| Kinematic Viscosity ($\nu$) | 1 | 1 $\mu\text{m}^2/\mu\text{s}$ |
| Time step ($dt$) | 0.125 | 0.125 $\mu\text{s}$ |
| The density of carrier fluid ($\rho$) | 1025 | 1025 $\text{fg}/\mu\text{m}^3$ |
| Bending Rigidity of RBC membrane ($\kappa$) | 1 | 1 pN $\mu\text{m}$ |
| Membrane Elasticity according to Skalak's Law (B, C, D) | B = 5, C = 5000, D = 35 | B = 5 pN/$\mu\text{m}$, C = 5000 pN/$\mu\text{m}$, D = 35 pN/$\mu\text{m}$ |
| Rayleigh Damping parameters for modelling viscosity ($\alpha_D$, $\beta_D$) | $\alpha_D = 0$, $\beta_D = 0.01$ | $\alpha_D = 0$, $\beta_D = 0.01$ |
| PBD Damping ($\kappa_{\text{damping}}$) | $\kappa_{\text{damping}} = 0.6$ | $\kappa_{\text{damping}} = 0.6$ |
| Volume Conservation constraint ($k_{\text{volume}}$) | $k_{\text{volume}} = 250$ | $k_{\text{volume}} = 250$ |

## Notes

### Competing Interest Statement

The authors have declared no competing interest.

