## Supplementary Information for "Dynamics of a Lipid Vesicle across a Microfluidic Constriction: How does the Fluidity of the Encapsulation and its Micro-Environment Matter?"

### 1. Vesicle dynamics obtained from 3D simulations

This section describes the different dynamics experienced by a singled-out vesicle as it passes through a constricted microfluidic pathway**.** Figure S1 (a-c) depicts the stretching, rolling, and tumbling dynamics, respectively, as characterized by the evolution of the orientation angle as a function of axial displacement. As mentioned earlier in section 3.1 in the main text, during stretching, vesicles undergo minimal changes in orientation angle ($\theta \approx 0$)  (Fig. S1(a), S2 (a)). In contrast, tumbling motion (Fig. S2(c), Fig. S1(c)) shows a drastic change in $\theta$, including a discontinuity from $90^o$ to $-90^o$. On the other hand, rolling dynamics (Fig. S2(b); Fig. S1 (b)) exhibit periodic variation in $\theta$ upon rotation about an axis orthogonal to the direction of flow. The insets in Figure S1 (a-c) depict the shape deformation of vesicles hallmarked by a parameter called deformation index ($DI=a-b/a+b$); where $a,b$ are the half major and minor axis lengths of a nearly elliptic vesicle under consideration.

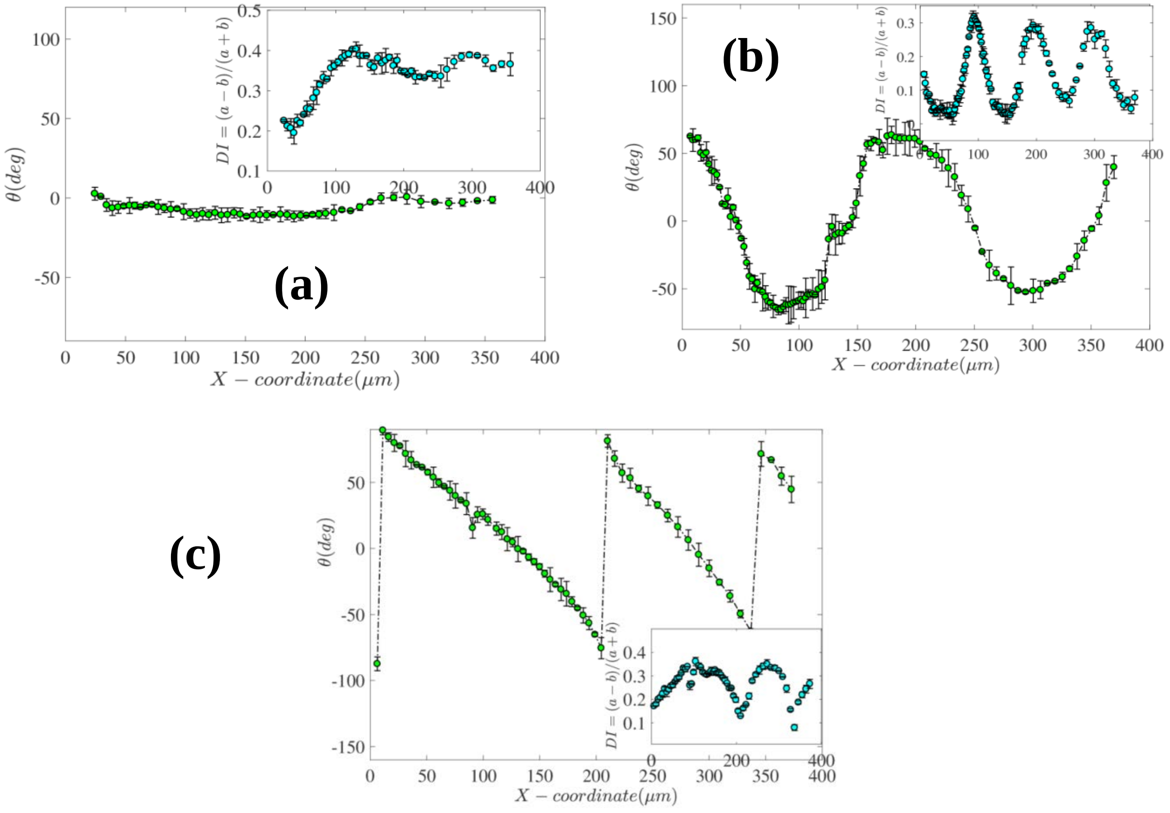


**Figure S1.** Representative behavior of an individual vesicle (**ν**=0.98) undergoing three different dynamics, namely stretching (a), rolling (b), and tumbling (c) motion while entering a microfluidic constriction obtained from 3D simulations. Insets of Fig. S1 (a to c) show the shape deformation (deviation from initial shape) manifested by the Deformation Index (*DI*) as a function of downstream position($X$) in the channel.

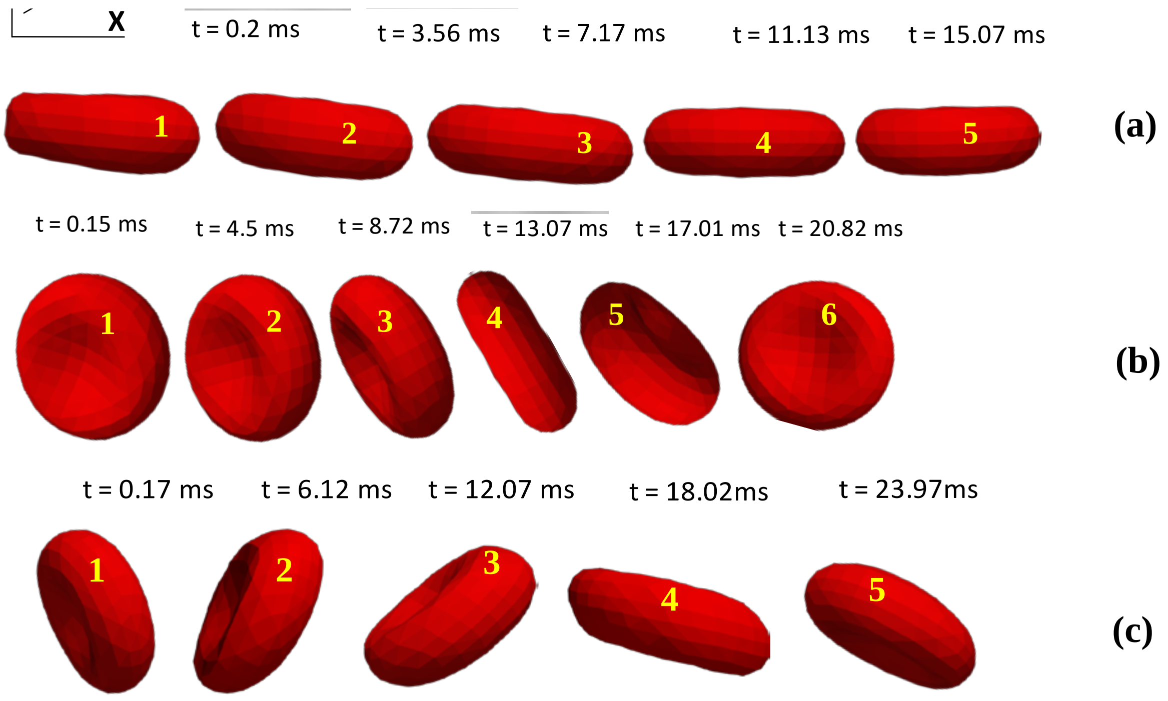


**Figure S2.** Time-lapse images of an individual vesicle (**v**=0.98) undergoing 3 different dynamics, namely stretching (a), rolling (b), and tumbling(c), respectively while entering a microfluidic constriction obtained from 3D numerical simulations.

### 2. Membrane bending modulus estimation from inverse mapping

The inverse mapping method has been used to estimate the value of membrane bending modulus by coupling the numerical and experimental data for a fixed identity vesicle undergoing stretching dynamics while migrating through the tapered region prior to the constriction. The simulations are carried out initially with a guess value of $E_B$, and the findings from the same in terms of stretch ratio vs. x-position (inset in Figure S3.) was superimposed with the experimental observation for the similar parametric space for different Glutaraldehyde concentration. An error minimization technique was employed to obtain the desired $E_B$ value resulting in a collapse of experiment and simulation results. The findings are represented in a tabular form (Table S1).

***Table S1. Membrane bending modulus as a function of Glutaraldehyde concentration***

| Gluta. Concentration (% v/v) | Max. stretch ratio | Normalized bending modulus |
|---|---|---|
| 0 | 1.315 | 0.002 |
| 5 | 1.236 | 0.053 |
| 10 | 1.2 | 0.084 |

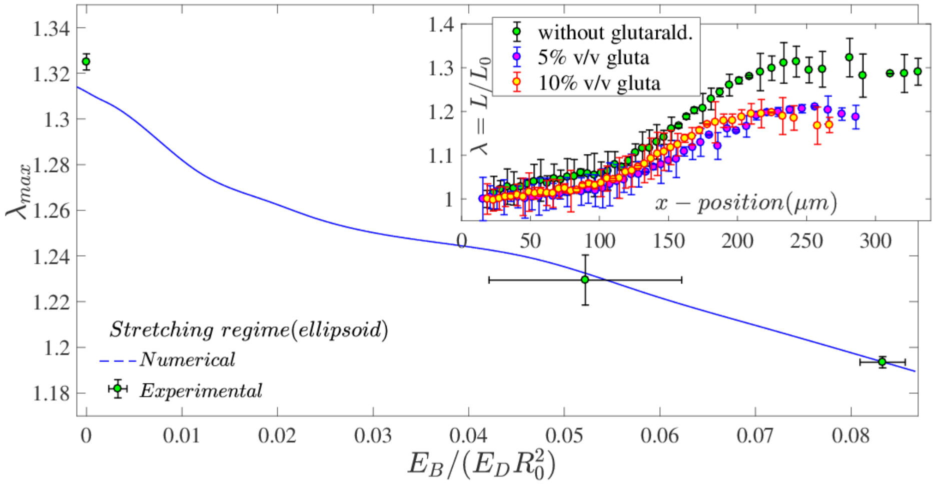


**Figure S3.** Variation of the maximum value of stretching ratio as a function of normalized membrane bending rigidity modulus for a vesicle undergoing stretching. Inset depicts the evolution of stretch ratio as a function of axial position for different Glutaraldehyde concentrations. It is evident from the inset figure that, with an increase in glutaraldehyde concentrations, the stretching is gradually arrested due to the increment of the bending modulus of the vesicle membrane. The open symbols in the figure demarcate the experimental findings, while the solid lines represent the numerical simulation results.

### 3. Size distribution of GUV samples obtained from electroformation

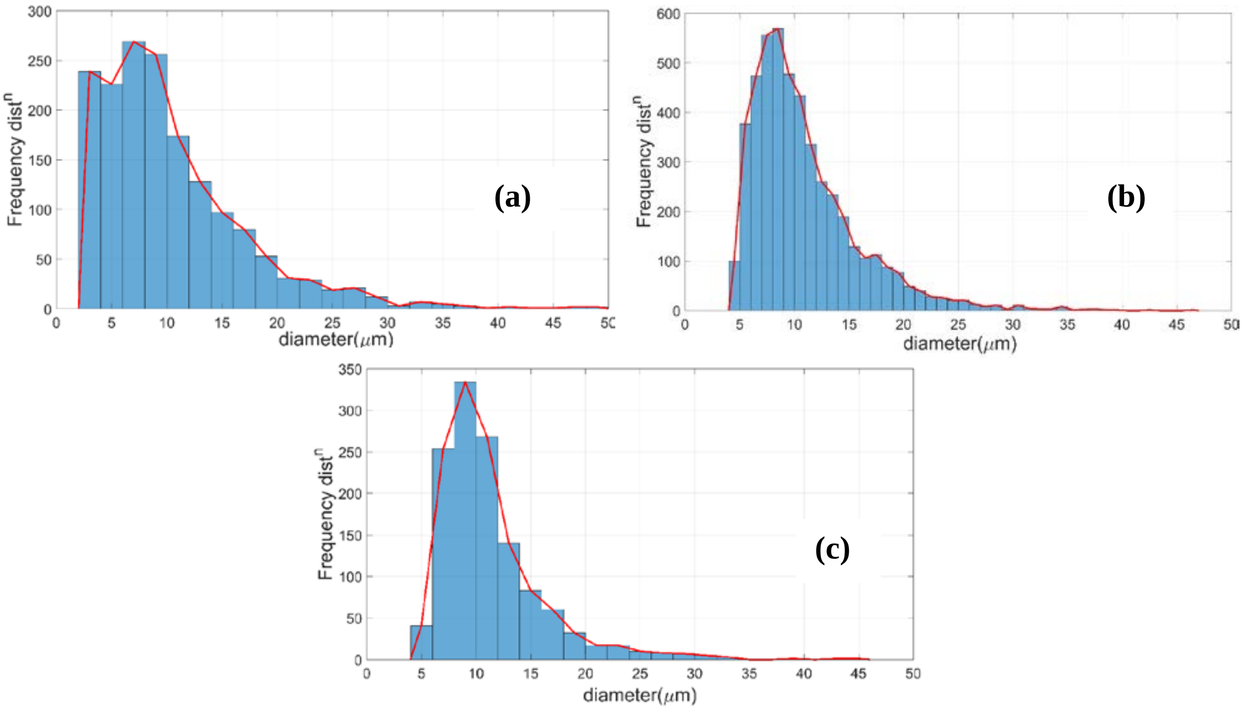


**Figure S4.** Size distribution of GUV samples obtained from electroformation: The histogram represents the variation in vesicle effective diameter ($2R_0$) as a function of its frequency/number obtained from sampling upon (a) 1650 nos. (b) 4605 nos. and, (c) 1261 nos. samples on different days.

#### Supporting video captions

**Movie S1:** A giant lipid-vesicle undergoing stretching dynamics (ellipsoid shape transition) while passing through the tapered region before entering the constriction under Poiseuille flow.

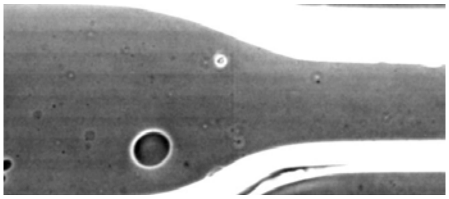

**Movie S2:** 2D-BEM simulation of an eccentrically placed vesicle undergoing stretching motion while moving through the tapered region.

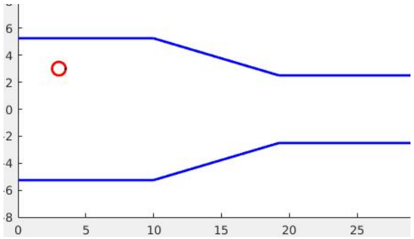


**Movie S3** Experimentally observed rolling motion of an eccentrically placed vesicle while passing through the tapered region of the rectangular cross-section microchannel.

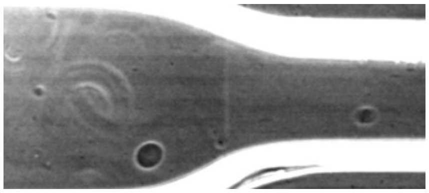

**Movie S4** 2D-BEM simulation of an eccentrically placed vesicle, undergoing rolling motion while moving through the tapered region.

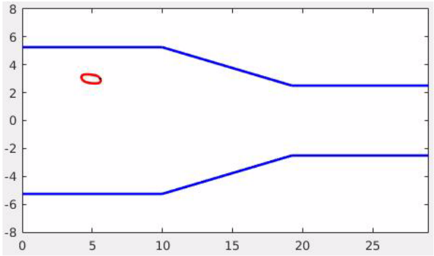


**Movie S5** Experimentally observed the tumbling motion of an eccentrically placed equiviscous ($\eta_r$ ~1) vesicle while passing through the tapered region of the rectangular microchannel.

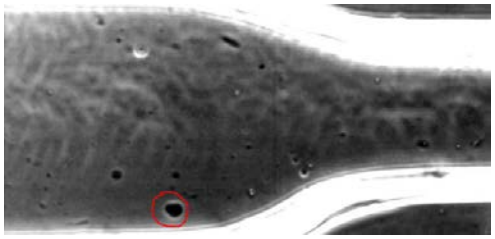

**Movie S6** 2D-BEM simulation of an eccentrically placed undergoing Tumbling motion while moving through the tapered region.

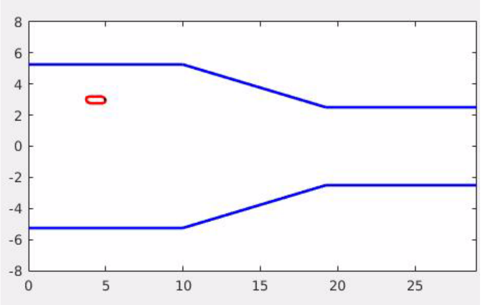


**Movies S7-S9:** 3D simulations of vesicle dynamics through constriction.
